# A new mechanism for a familiar mutation – bovine *DGAT1* K232A modulates gene expression through multi-junction exon splice enhancement

**DOI:** 10.1101/2020.02.04.934562

**Authors:** Tania Fink, Thomas J Lopdell, Kathryn Tiplady, Renee Handley, Thomas JJ Johnson, Richard J Spelman, Stephen R Davis, Russell G Snell, Mathew D Littlejohn

## Abstract

The *DGAT1* gene encodes an enzyme responsible for catalysing the terminal reaction in mammary triglyceride synthesis, and underpins a well-known pleiotropic quantitative trait locus (QTL) with a large influence on milk composition phenotypes. Since first described over 15 years ago, a protein-coding variant K232A has been assumed as the causative variant underlying these effects, following *in-vitro* studies that demonstrated differing levels of triglyceride synthesis between the two protein isoforms. In the current study, we used a large RNAseq dataset to re-examine the underlying mechanisms of this large milk production QTL, and hereby report novel expression-based functions of the chr14 g.1802265AA>GC variant that encodes the *DGAT1* K232A substitution. Using expression QTL (eQTL) mapping, we demonstrate a highly-significant mammary eQTL for *DGAT1*, where the K232A mutation appears as one of the top associated variants for this effect. By conducting *in vitro* expression and splicing experiments in bovine mammary cell culture, we further show modulation of splicing efficiency by this mutation, likely through disruption of an exon splice enhancer as a consequence of the allele encoding the 232A variant. Although the relative contributions of the enzymatic and transcription-based mechanisms now attributed to K232A remain unclear, these results suggest that transcriptional impacts contribute to the diversity of lactation effects observed at this locus.

## Introduction

A lysine to alanine amino acid substitution (K232A) encoded by a mutation in the diacylglyercol O-acyltransferase 1 (*DGAT1*) gene has major impacts on bovine lactation traits, the most substantial being its impact on milk fat percentage [1,2]. This substitution results from an AA to GC dinucleotide substitution in exon eight of *DGAT1*, and likely constitutes the most widely studied and validated variant in association analyses of bovine milk characteristics (initially described by Grisart et al (2002), with >1000 Google Scholar citations to date). The *DGAT1* gene encodes an enzyme responsible for catalysing the terminal reaction in the mammary triglyceride synthesis pathway [3], and the *DGAT1* K allele has been shown [4] to synthesise more triglycerides *in vitro* when compared to the A allele. Aside from the *DGAT1* K232A mutation, an additional polymorphism 5′ of the transcription start site of the gene has also been shown to associate with milk fat percentage [5]. This variant, a variable number tandem repeat (VNTR) expansion, was speculated to play a role in bovine milk composition by increasing the number of putative transcription factor binding sites [5]. However, functional testing of the VNTR variant did not show any differences in *DGAT1* expression between QTL genotypes in cell culture [6]. This finding largely put the competing, gene expression-based hypothesis of the *DGAT1* milk fat effect to rest, with enzymatic differences deriving from the K232A mutation now widely assumed as the underlying mechanism.

Since these initial analyses >10 years ago, further functional characterisation of the K232A mutation has been largely absent. Having generated a large, mammary RNAseq dataset, however, we recently had the opportunity to re-examine this locus for potential regulatory effects impacting *DGAT1*, among other loci [7], and demonstrated a strong *DGAT1* expression-QTL (eQTL) in the mammary gland. Importantly, the expression of *DGAT1* transcripts was associated with K232A genotype (and thus milk fat percentage). Based on this observation, we have investigated mechanisms by which the K232A variant might mediate this effect. For the first time, we present functional and statistical evidence suggesting splice-enhancement-based regulatory control of *DGAT1* transcripts as an explanation for the eQTL, and by inference the milk fat and lactation effects attributed to this gene and mutation.

## Results

### *DGAT1* K232A is strongly associated with *DGAT1* transcript abundance in the lactating mammary gland

To test for regulatory effects on *DGAT1* transcript levels, we performed eQTL mapping using a mammary RNAseq dataset representing 375 lactating cows (Methods). Association testing was conducted using transformed mammary *DGAT1* transcript counts and 115 SNPs from the BovineHD panel, representing a 1Mbp interval of variants centred on (and also including) the K232A mutation. This analysis revealed a highly significant eQTL for *DGAT1*. Curiously, K232A was one of the top associated variants (P=1.59×10^−25^; Figure 1A), explaining 29.7% of the phenotypic variance in mammary *DGAT1* expression (Table 1).

**Figure 1.**
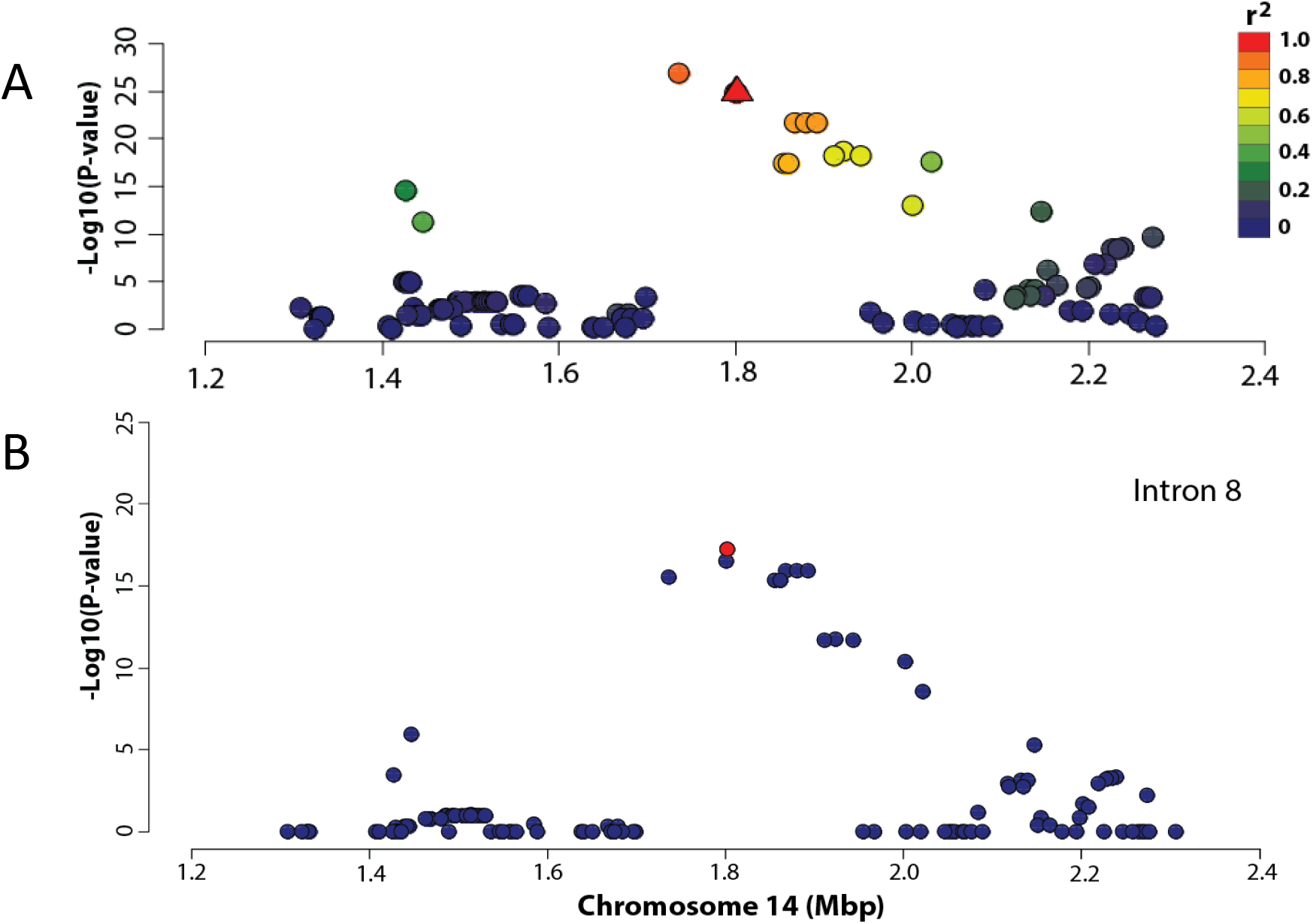
Expression QTL analysis at the *DGAT1* locus in bovine lactating mammary gland. A) The X-axis shows the position on chromosome 14 (millions of base pairs on the UMD3.1 reference genome), the Y-axis shows −log_10_ P-values of marker association for the 115 SNPs from the BovineHD panel (plus K232A) in the 1 Mbp interval centred on *DGAT1* K232A (denoted as a triangle). B) Marker association with intron 8 splicing efficiency for the 116 SNPs in the 1 Mbp interval. The K232A marker is coloured red.

**Table 1.**
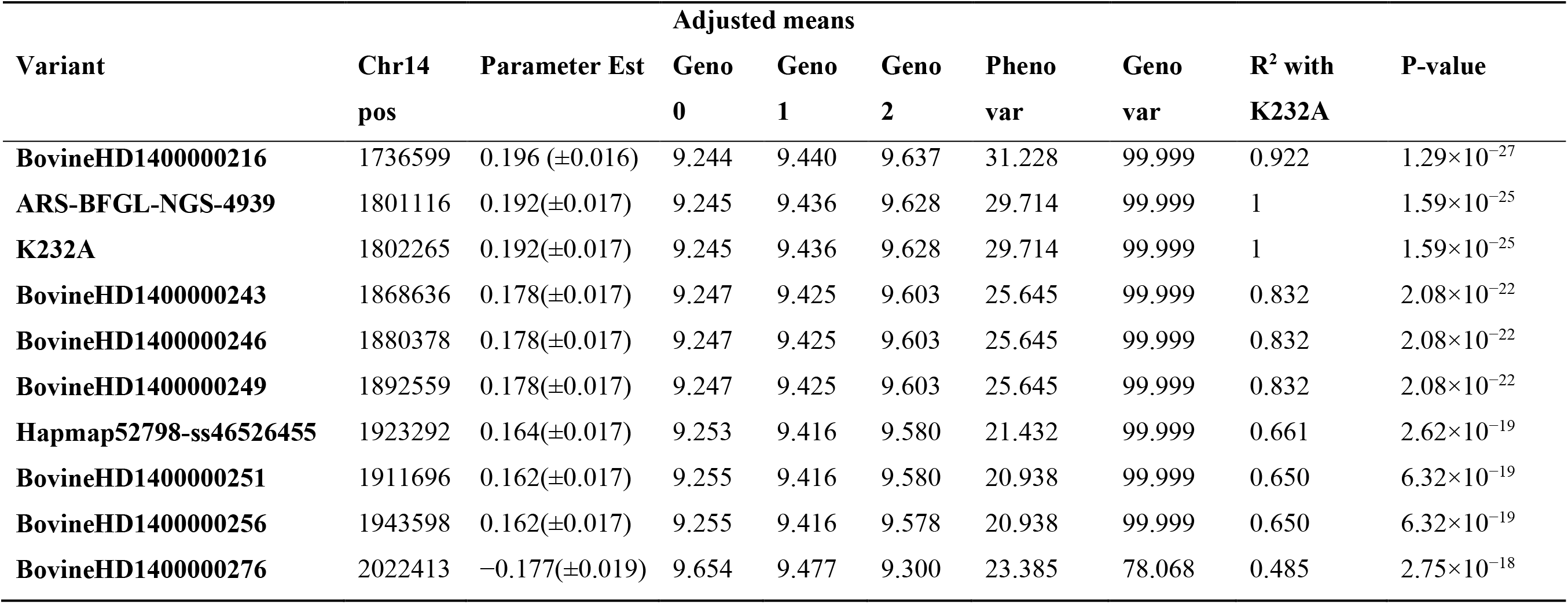
Mammary *DGAT1* expression association statistics for the top 10 BovineHD variants. The positions of these SNP variants on chromosome 14 are relevant to the UMD3.1 reference. Parameter estimates (±SE) and parameter-adjusted means are shown in units of VST-transformed RNAseq read counts. The linkage disequilibrium R^2^ values for each SNP are shown relative to the K232A variant.

While K232A was significantly associated with *DGAT1* expression, the most highly associated marker for this signal was BovineHD1400000216, which is located approximately 59 kbp upstream of *DGAT1* at chr14: 1736599 (P=1.29×10^−27^; Figure 1A). This marker is in strong LD with K232A, with an R^2^ value of 0.92 (Table 1). Notably, the milk fat percentage-increasing K allele was the same allele associated with increased *DGAT1* expression in this analysis. Those animals homozygous for the K allele had a mean transformed read count for *DGAT1* of 9.628 (±0.024), whereas for those animals with the A allele this value was 9.245 (±0.026). Heterozygous animals had a mean transformed read count of 9.436 (±0.019), intermediate between the two opposing homozygous classes (Table 1). The frequency of the *DGAT1* K allele was 0.51 in the RNAseq population.

### *DGAT1* K232A associates with alternative splicing of *DGAT1* exon 8

Based on a previous report of the influence of *DGAT1* K232A on the generation of an alternative *DGAT1* splicing isoform [4], we wondered whether this alternative splicing of exon 8 might give rise to the observed eQTL in the RNAseq dataset. The relative usage of the reference and alternative exons was quantified using DEXSeq [8], to create a phenotype for the *DGAT1* exon 8 alternative splicing ratio (see Methods). Analysis of the splicing ratio phenotype in conjunction with K232A genotype, revealed a significant difference in the ratio of reads splicing at the alternative splice site and the constituent splice site for exon 8 based on K232A genotype (P=4.60×10^−6^).

Notably, the *DGAT1* expression-increasing K allele was the same allele associated with increased alternative splicing of *DGAT1* exon 8 (Table 2). However, this is unlikely to explain the eQTL, as the alternative isoform differs through the ‘intronification’ [4] of the majority of exon 8, and is therefore expected to result in a reduction in the number of reads mapping to the exons of the gene overall.

**Table 2.**
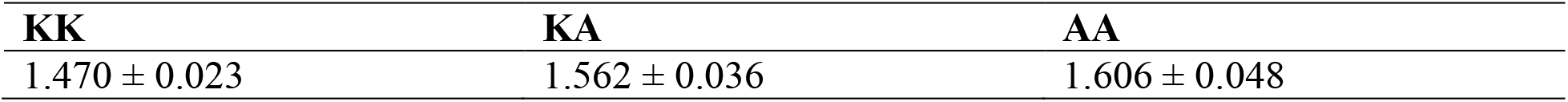
Alternative splicing ratios for *DGAT1* exon 8. Columns indicate genotypes for the K232A variant. Splicing ratios are shown ±standard errors.

### *DGAT1* K232A disrupts a putative consensus exon splice enhancer

Given the strong statistical association of the K232A variant with mammary *DGAT1* expression and alternative isoform abundance, we wondered whether the variant might be associated with other splicing effects. The mutation sits within 15 bp of the intron/exon boundary, highlighting other splicing-based mechanisms as a possible explanation for the observed eQTL. To examine this possibility, the genomic sequence surrounding K232A was examined for the presence of exon splice enhancer (ESE) motifs. The RESCUE-ESE analysis tool [9] was used to annotate *DGAT1* exon 8 for predicted ESEs. For both *DGAT1* alleles, the first 23 nucleotides of the 5′ end *DGAT1* exon 8 were examined using the default 238 human ESE panel. This analysis revealed the presence of two predicted ESE motifs in the 5′ end of *DGAT1* exon 8 which both overlap K232A (Figure 2).

**Figure 2.**
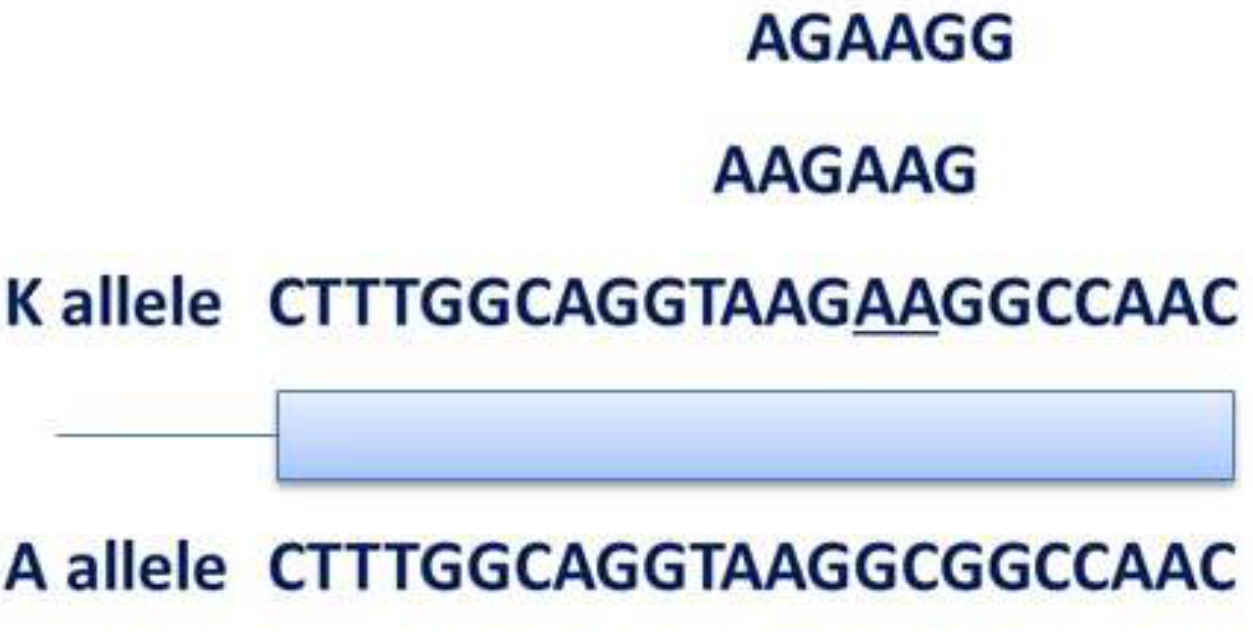
Schematic of the 5′ end of DGAT1 exon 8 with ESE motifs overlapping the K232A amino acid substitution. The AA>GC MNP responsible for the K232A substitution is underlined in the DGAT1 K allele.

Notably, these ESE motifs (AG<u>AA</u>GG and AAG<u>AA</u>G) were located at chr14: 1802263–1802268 and chr14: 1802262–1802267 (UMD3.1 genome build) respectively, and were only encoded by the K allele. These ESEs were disrupted by the AA>GC MNP (Figure 2). As such, this polymorphism could be expected to disrupt these two ESE motifs when the 232A allele is present (GC), which is the same allele associated with decreased mammary *DGAT1* expression and milk fat percentage.

### *DGAT1* K232A associates with splicing efficiency of *DGAT1* introns

Having observed differential expression of *DGAT1* and the possible disruption of ESEs based on K232A genotype, we next investigated whether this variant associated with the efficiency of splicing at the neighbouring splicing junction. To this end, a splicing efficiency phenotype was derived for intron 8, which involved quantification of the percentage of total *DGAT1* RNAseq reads that mapped to this intron junction (see Methods).

Association analysis between the percentage of RNAseq reads mapping to intron 8 and K232A and the 115 SNPs from the BovineHD panel revealed a strong splicing efficiency effect, with K232A the most significantly associated variant (P=6.29×10^−18^, Figure 1B). Importantly, the direction of effect for the splicing efficiency effect was consistent with a mechanism that might explain the *DGAT1* eQTL, such that the K allele was associated with an increased percentage of completely spliced transcripts at the exon 8 junction. Of the three genotype classes, the KK animals had the highest percentage of completely spliced transcripts with only 1.16% of *DGAT1* RNAseq reads mapping to intron 8, while the AA animals had the lowest percentage of completely spliced transcripts with 1.94% of reads mapping to intron 8. Heterozygous AK animals showed intermediate levels of splicing (1.53% of reads mapping to intron 8). As the K allele is also associated with increased *DGAT1* expression, this suggested splicing efficiency as a potential limiting mechanism for the production of fully spliced mRNA.

Given the observation of differential splicing efficiency of intron 8, we conducted association analyses for each of the other 13 *DGAT1* junctions, using the same subset of 116 markers. This revealed significant associations at eight additional *DGAT1* junctions (Table 3), with the most significant impacting the intron 2 junction (P=5.25×10^−46^). For the nine significantly impacted junctions, K232A was the lead SNP for the association at three junctions (introns 2, 7 and 8). Interestingly, many of these junctions are physically distant to the K232A variant and the proposed ESE motif. For example, in base position terms, intron 2 is several kb from K232A yet was the most significant splicing efficiency effect in this analysis (Table 3).

**Table 3.**
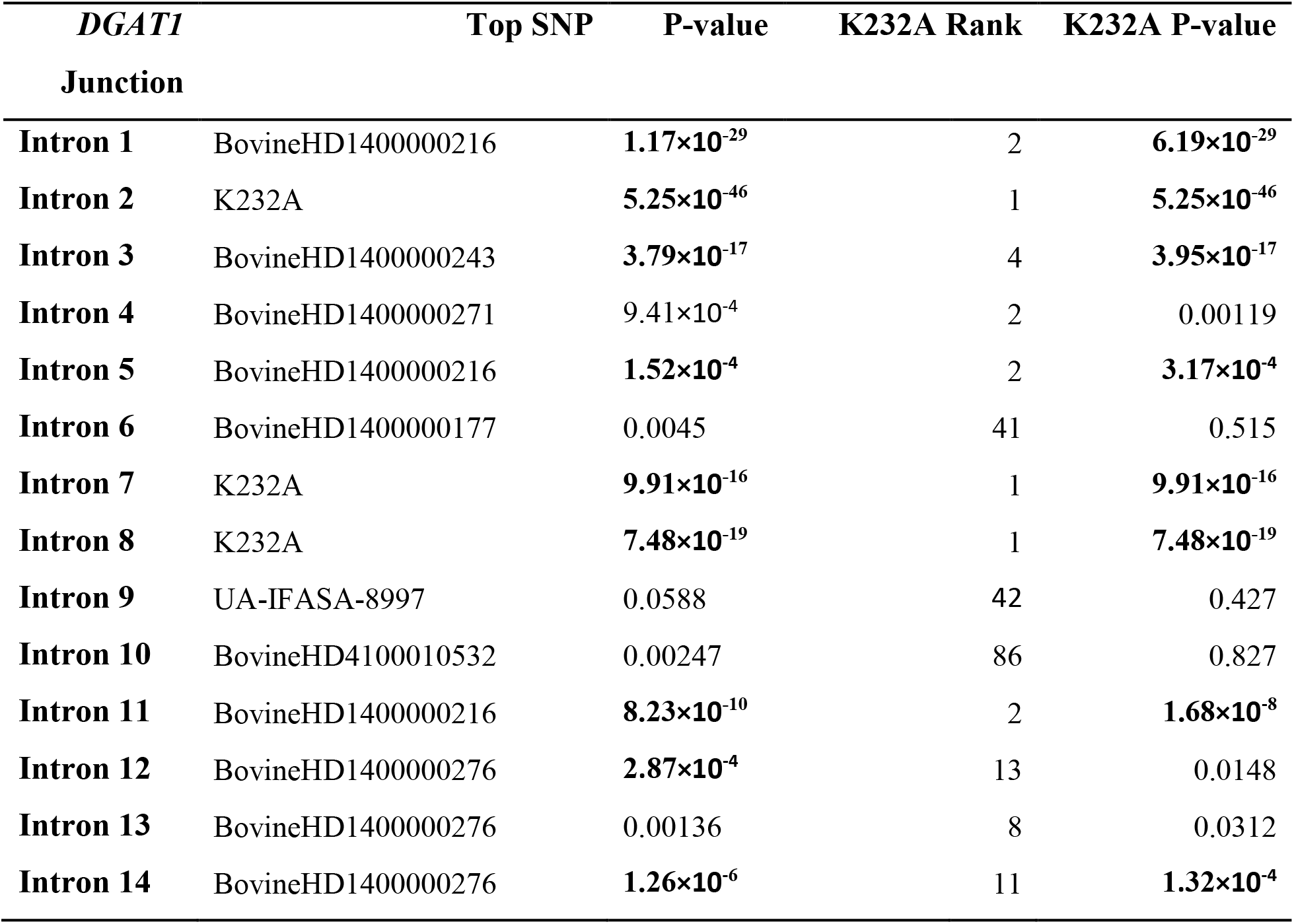
*DGAT1* junction splicing efficiency association statistics for the top BovineHD panel variants and K232A. The fourth column displays the ranking of the association between K232A and splicing at the indicated junction, compared to all the markers. The significance of the K232A association is presented in the fifth column, compared to the P-value of the most-significantly associated variant in the third column (identical if the most significant variant was K232A). P-values lower than the Bonferroni threshold (P=4.31×10^−4^) are highlighted using bold type.

### *DGAT1* K232A alters *DGAT1* splicing efficiency *in vitro*

While *DGAT1* K232A is associated with the expression of *DGAT1* and splicing efficiency of multiple *DGAT1* junctions, the observation of a higher-ranking SNP-chip marker in the eQTL analysis presents an alternative, equally plausible hypothesis that an unknown promoter or other *cis*-regulatory variant drives the gene expression effect. To test the possibility of K232A simply being in linkage disequilibrium (LD) with another regulatory mutation, cell-based experiments were undertaken to remove the two K232A transcript isoforms from their genomic context.

For these *in vitro* expression experiments, plasmid-based mini-gene constructs were synthesised and cloned into a pcDNA3.1 vector backbone under the control of a CMV promoter (Supplementary Figure 1). These constructs were designed to represent the identical intron-exon structure of genomic *DGAT1* with the exception that introns 1 and 2 had been removed (due to plasmid size constraints, see Methods). The resultant expression vectors differed only by the AA and GC alleles that encode the K/A residues, allowing spliced and unspliced transcripts to be compared between constructs. Quantification of splicing was achieved by qPCR, targeting four individual intron/exon boundaries that either showed (introns 3 and 7), or did not show (introns 5 and 13) major differential splicing efficiency effects *in vivo* (Table 3; Supplementary Figure 1).

Following transfection of the *DGAT1* constructs into the bovine mammary cell line MAC-T [29], the alternate mini-gene alleles were found to recapitulate the splicing effects observed in the lactating mammary dataset. Transcripts generated from cells expressing the K allele showed higher splicing ratios at the intron 3 and intron 7 junctions compared with those cells transfected with the A allele (P=9.37×10^−4^ and P=9.05×10^−11^, respectively; Table 4). For the intron 5 and 13 junctions, there were no significant differences in splicing efficiency (P=0.256 and P=0.497, respectively; Table 4). At the intron 7 junction, analysis of the mean expression of spliced transcripts showed increases in expression in cells transfected with the K allele (P=0.009; Figure 3). Although the other three junctions also showed numerical increases in the expression of spliced transcripts for the K construct, these effects were non-significant after adjustment for multiple hypothesis testing (Table 4; Bonferroni threshold P=0.0125). None of the junctions showed significant differences in the mean expression of unspliced transcripts between constructs. As such, the increased expression of spliced mRNAs appeared to derive from post-transcriptional mechanisms, likely as a consequence of enhanced splicing efficiency, at least as assayed at the intron 7 junction that was significant for these effects.

**Figure 3:**
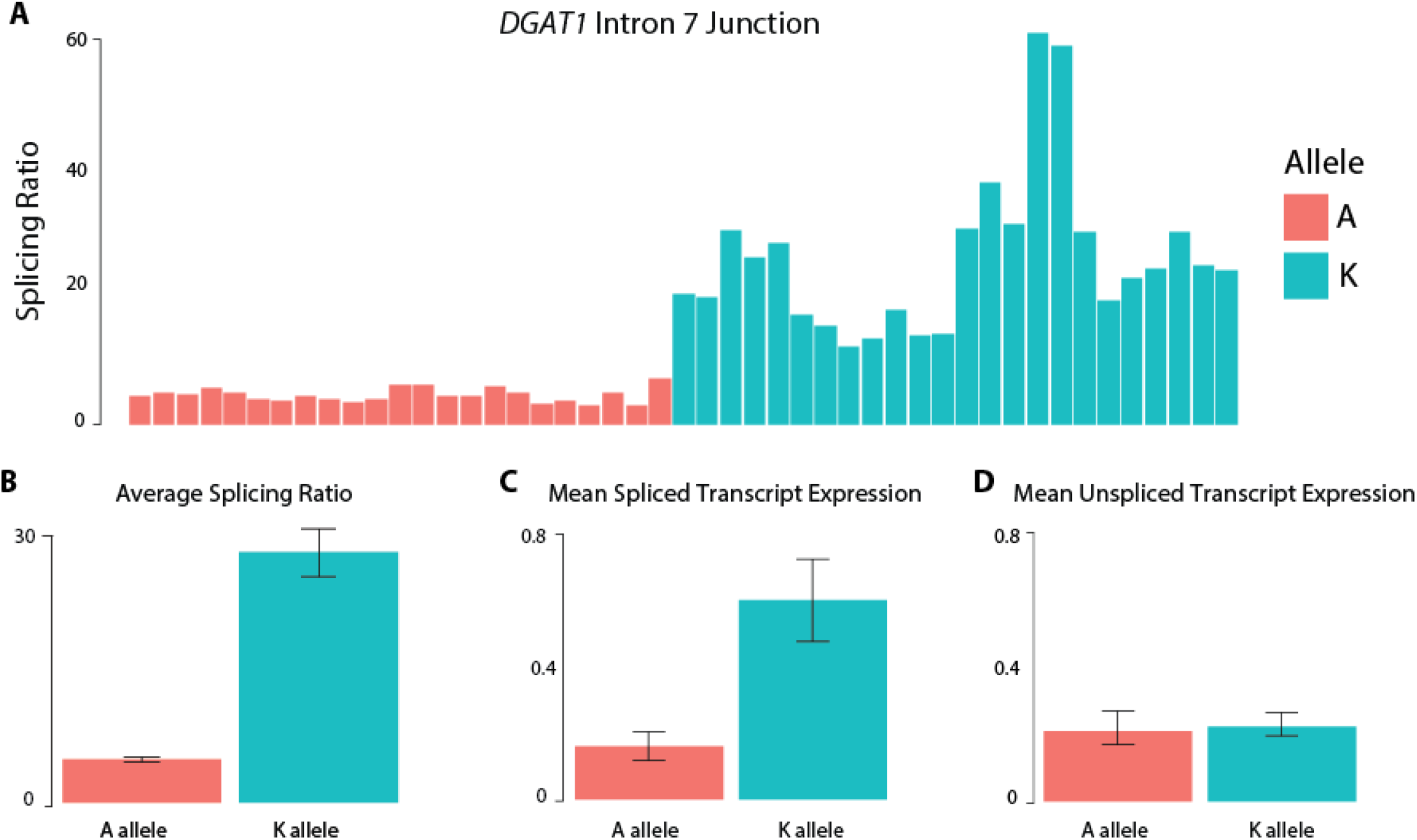
Cell-based functional testing of DGAT1 K232A influence on splicing efficiency at the DGAT1 intron 7 junction. Figure A represents the splicing ratio (spliced transcripts:unspliced transcripts) measured by qPCR in each of the individual replicates for the intron 7 junction. Figure B represents the average splicing ratio for the intron 7 junction for the two DGAT1 K232A alleles. The error bars represent the standard deviation across all samples. Figures C and D represent the mean spliced and unspliced transcripts for the intron 7 junction, respectively. The error bars represent the standard error of the difference between means.

**Table 4.** **Measurements of splicing efficiency** at *DGAT1* intron 3, 5, 7 and 13 junctions in mammary cell culture for the 232K and 232A *DGAT1* plasmids. Bonferroni threshold P=0.0125.

| Measurement | DGAT1 Junction | K allele | A allele | P-value |
| --- | --- | --- | --- | --- |
| Splicing ratio | 3 | 0.253 ($\pm 0.009$) | 0.173 ($\pm 0.005$) | **$9.37 \times 10^{-4}$** |
| | 5 | 143.18 ($\pm 10.23$) | 133.52 ($\pm 9.68$) | 0.256 |
| | 7 | 28.49 ($\pm 2.77$) | 5.55 ($\pm 0.74$) | **$9.05 \times 10^{-11}$** |
| | 13 | 92.90 ($\pm 5.86$) | 90.92 ($\pm 5.83$) | 0.411 |
| Spliced mRNA expression | 3 | 0.111 ($\pm 0.002$) | 0.082 ($\pm 0.002$) | 0.015 |
| | 5 | 0.946 ($\pm 0.3238$) | 0.421 ($\pm 0.160$) | 0.097 |
| | 7 | 0.545 ($\pm 0.117$) | 0.255 ($\pm 0.047$) | **0.009** |
| | 13 | 0.984 ($\pm 0.219$) | 0.549 ($\pm 0.233$) | 0.022 |
| Unspliced mRNA expression | 3 | 0.092 ($\pm 0.012$) | 0.0842 ($\pm 0.227$) | 0.376 |
| | 5 | 0.286 ($\pm 0.0415$) | 0.180 ($\pm 0.0350$) | 0.069 |
| | 7 | 0.216 ($\pm 0.030$) | 0.234 ($\pm 0.059$) | 0.383 |
| | 13 | 0.821 ($\pm 0.180$) | 0.503 ($\pm 0.124$) | 0.035 |

### *DGAT1* eQTL analysis at sequence resolution

Given the demonstration that K232A was associated with splice enhancement and mean mRNA expression *in vitro*, we re-visited association mapping of the locus using imputed whole genome sequence (WGS) data. In this analysis, we aimed to assess the relative contribution of the K232A variant to the *DGAT1* eQTL in the context of full sequence information, examining whether other sequence variants were substantially more associated with expression, or whether the K232A wholly explained this effect. Association analysis was conducted using 3128 genetic markers imputed from a WGS dataset that has been described elsewhere [10,11]. As for analysis of variants representing the BovineHD SNP-chip, this 1 Mbp interval was centred on the *DGAT1* K232A mutation, and analysed in conjunction with VST-transformed exonic *DGAT1* expression values. Like the previous association analysis using the BovineHD markers, this analysis revealed a strong eQTL for *DGAT1* in the mammary gland, with K232A remaining one of the top associated variants (P=1.59×10^−25^; Figure 4A).

**Figure 4:**
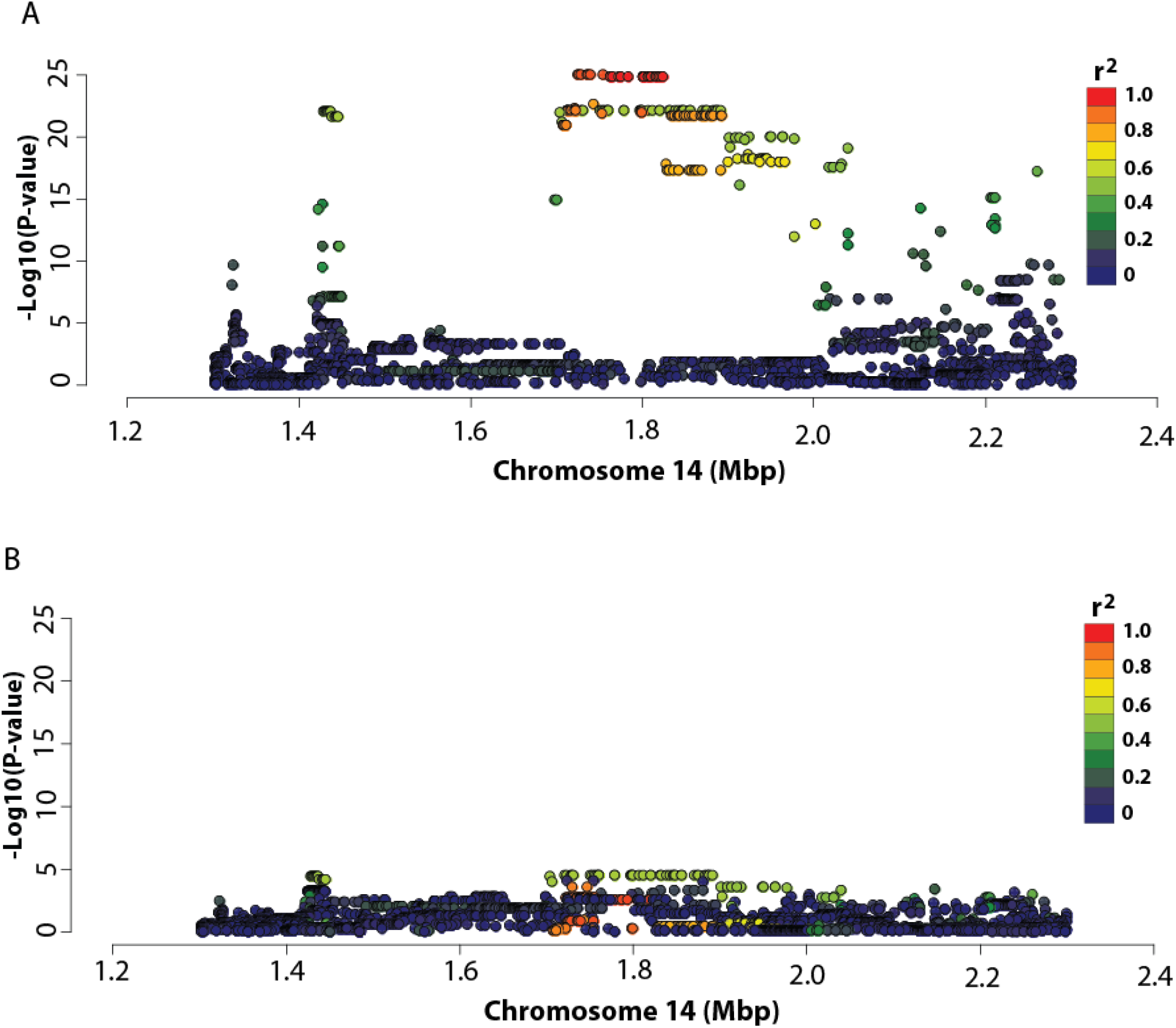
Expression QTL analysis at the *DGAT1* locus in bovine lactating mammary gland. Figure A shows the Manhattan plot for DGAT1 expression at the DGAT1 locus in the RNAseq animals (n=375). The X-axis shows chromosome 14 position (million base pairs); the Y-axis shows −log_10_ P-values of marker association for the 3,218 WGS-derived SNPs in the 1 Mbp interval centred on DGAT1 K232A. Figure B represents the Manhattan plot for DGAT1 expression at the DGAT1 locus in the RNAseq animals conditioned on K232A. Markers are coloured based on their correlations (R^2^) with K232A in both Figures A and B.

The most highly associated markers for this signal were rs209328075 and rs209929366, located upstream of *DGAT1* at chr14:1730455 and chr14:1747132, respectively (P=2.31×10^−28^; Table 5). These markers exhibited identical association statistics with mammary *DGAT1* expression. They were also highly correlated with K232A, exhibiting an R^2^ value of 0.88. When K232A was fitted as a covariate in the association model, the association of the two lead variants was greatly reduced (Figure 4B). These associations were non-significant when applying a Bonferroni correction, though significant in the absence of this correction (P=0.000263; Bonferroni threshold P=1.60×10^−5^). Of greater note, a cluster of 39 variants in perfect linkage disequilibrium (LD) with each other, but only modestly correlated with K232A (R^2^=0.548) were significant (P=3.58×10^−5^) in these models. These variants explained 10.32% of the residual phenotypic variance in mammary *DGAT1* expression, suggesting the possibility of another, functionally independent regulatory effect at the locus. Interestingly, some of the markers most highly associated with the residual *DGAT1* eQTL signal reside several kb upstream of the transcription start site of the gene (chr14:1428907–1754446; Table 6), representing candidate variants that may impact an additional upstream promoter or other regulatory feature of *DGAT1*.

**Table 5.** Mammary *DGAT1* expression association statistics for top WGS-derived variants. The positions of these SNP variants are indicated, with parameter estimates shown with standard errors in units of VST-transformed RNAseq read counts. The genetic and phenotypic variance explained by each SNP, along with parameter-adjusted means for each of the three genotypes classes is indicated. The linkage disequilibrium R^2^ values for each SNP relative to the *DGAT1* K232A variant is shown, with the P-values indicated in the right most column.

| Variant | Chr14<br>pos | Parameter Est | Adjusted means | | | Pheno<br>var | Geno<br>var | $R^2$ with<br>K232A | P-value |
| --- | --- | --- | --- | --- | --- | --- | --- | --- | --- |
|  |  |  | Geno<br>0 | Geno<br>1 | Geno<br>2 |  |  |  |  |
| **rs209328075** | 1730455 | 0.1927($\pm 0.0161$) | 9.251 | 9.446 | 9.642 | 31.32 | 99.99 | 0.881 | $2.31 \times 10^{-28}$ |
| **rs209929366** | 1747132 | 0.1927($\pm 0.0161$) | 9.251 | 9.446 | 9.642 | 31.32 | 99.99 | 0.881 | $2.31 \times 10^{-28}$ |
| **rs208091850\*** | 1722033 | 0.1961($\pm 0.0164$) | 9.244 | 9.440 | 9.637 | 31.23 | 99.99 | 0.922 | $1.29 \times 10^{-27}$ |
| **rs208417762^** | 1756075 | 0.1969($\pm 0.0166$) | 9.240 | 9.437 | 9.634 | 31.33 | 99.99 | 0.968 | $2.38 \times 10^{-27}$ |
| **rs135458711+** | 1724688 | 0.1906($\pm 0.0167$) | 9.250 | 9.440 | 9.631 | 29.46 | 99.99 | 0.952 | $1.10 \times 10^{-25}$ |
| **K232A&** | 1802265 | 0.1919($\pm 0.0169$) | 9.244 | 9.436 | 9.628 | 29.71 | 99.99 | 1 | $1.59 \times 10^{-25}$ |

\*31, ^27, +7 and &20 additional genetic variants, respectively had the same association signal for *DGAT1*. These variants were statistically indistinguishable from each other and are not included in this table in the interest of size.

**Table 6.** Mammary *DGAT1* expression association statistics for top sequence variants conditioned on *DGAT1* K232A. The positions of these SNP variants are indicated, with parameter estimates shown with standard errors in units of VST-transformed RNAseq read counts. The genetic and phenotypic variance explained by each SNP, along with parameter-adjusted means for each of the three genotypes classes is indicated. The linkage disequilibrium R^2^ values for each SNP relative to the *DGAT1* K232A variant is shown, with the P-values indicated in the right most column.

| Variant | Chr14<br>pos | Parameter Est | Adjusted means |  |  | Pheno<br>var | Geno<br>var | R<sup>2</sup> with<br>K232A | P-value |
| --- | --- | --- | --- | --- | --- | --- | --- | --- | --- |
|  |  |  | Geno<br>0 | Geno<br>1 | Geno<br>2 |  |  |  |  |
| **rs472613236\*** | 1721117 | 0.1013(±0.0242) | 9.361 | 9.463 | 9.564 | 10.32 | 99.99 | 0.548 | 3.58x10<sup>-5</sup> |
| **rs383105805^** | 1428907 | 0.1002(±0.0240) | 9.363 | 9.463 | 9.564 | 10.13 | 99.99 | 0.539 | 3.88x10<sup>-5</sup> |
| **rs109448144** | 1704351 | -0.1004(±0.0240) | 9.564 | 9.464 | 9.364 | 10.06 | 99.99 | 0.544 | 4.14x10<sup>-5</sup> |
| **rs137587412&** | 1438890 | 0.0975(±0.0243) | 9.366 | 9.463 | 9.561 | 9.62 | 99.99 | 0.546 | 7.08x10<sup>-5</sup> |
| **rs445906781** | 1723278 | 0.1165(±0.0295) | 9.424 | 9.540 | 9.657 | 4.95 | 99.99 | 0.090 | 9.71x10<sup>-5</sup> |
| **rs476272800** | 1754446 | 0.1165(±0.0295) | 9.424 | 9.540 | 9.657 | 4.88 | 99.99 | 0.853 | 9.71x10<sup>-5</sup> |

\*39, ^9 and &5 additional genetic variants, respectively had the same association signal for *DGAT1*. These variants were statistically indistinguishable from each other and are not included in this table in the interest of size.

## Discussion

A pleiotropic QTL with a large influence on milk composition resides on the centromeric end of bovine chromosome 14, underpinned by the *DGAT1* gene. Although there was initially speculation as to the specific genetic variant and mechanism responsible for the QTL [1,5], *in vitro* functional evidence showing that the two protein isoforms of *DGAT1* differed in their ability to synthesise triglycerides [4] led to the now-dominant hypothesis that the K232A amino acid substitution is the causative variant. In the current study, we sought to re-assess the potential role of transcriptional regulation as a mechanism underlying the *DGAT1* effects. To this end, we herein provide genetic and functional evidence as to how the *DGAT1* K232A mutation may influence milk composition through splice-site enhancement and consequent expression-based effects.

We had previously conducted association analysis using RNAseq-derived expression data and revealed a strong eQTL for *DGAT1* in lactating mammary tissue [7]. Importantly, the mammary *DGAT1 cis*-eQTL showed a similar genetic signal underpinning the milk production QTLs reported for this locus, with K232A highly associated with the gene expression effect. We have shown in the current study that animals bearing the K allele for *DGAT1* K232A possess greater mammary *DGAT1* expression compared to those animals bearing the A allele. This is of particular note given the K allele is the same allele associated with increased milk fat percentage[1], and more *DGAT1* enzyme as a consequence of increased mRNA could be expected to increase triglyceride synthesis.

The finding that the K232A variant is one of the most highly associated genetic markers with *DGAT1* expression is surprising, since previous *in vitro* investigations using RT-PCR found no difference in *DGAT1* mRNA expression based on the K232A genotype, albeit with limited numbers of animals [4] (N=24). Therefore, the effect of *DGAT1* K232A on milk fat production had been attributed to the enzymatic difference between the two DGAT1 isoforms, as the K allele had the greatest enzymatic activity *in vitro*, and associated with increased milk fat percentage [4]. Based on that previous study, it has been widely assumed that this enzymatic difference was the sole mechanism driving the effect of K232A on milk composition. In that same study, *DGAT1* K232A was shown to associate with an alternatively spliced transcript of *DGAT1* [4]. This isoform differs based on its utilisation of a splice site 6 bp upstream of K232A, resulting in the ‘intronification’ of the majority of exon 8. The protein encoded from this isoform is predicted to have an internal deletion of 22 amino acids, and is assumed to be non-functional based on its inability to synthesise triacylglycerides *in vitro* [4]. The proportion of this alternative isoform is approximately 10% of the total *DGAT1* transcripts, and it was shown in the same manuscript that first described this alternative transcript that the K allele results in an increase in its expression *in vitro* [4]. In line with this observation, the ratio of the alternative isoform to the full-length form differed by K232A genotype in the current dataset, with the animals bearing the K allele producing more of this isoform compared to those animals with the A allele.

Originally, we had hypothesised that the mammary *DGAT1* eQTL might be the result of increased alternative *DGAT1* isoform production. As the alternative isoform results in the intronification of the majority of exon 8, however, we view this as unlikely – since the K allele would present fewer sequence reads mapping to the standard exon definition, yet is the same allele associated with increased mean expression.

Regulatory control of gene expression is most commonly attributed to non-coding sequences; however, regulatory elements can also be part of the coding sequence, and coding variants, such as the dinucleotide substitution underlying *DGAT1* K232A, can influence gene expression through the modulation of auxiliary splicing elements. We hypothesised that the *DGAT1* K232A mutation may overlap one of these elements, and the data presented herein support this hypothesis. Use of the RESCUE-ESE tool to annotate the exonic sequence around K232A suggests two predicted ESE motifs (AGAAGG and AAGAAG) that overlap the K232A polymorphism, with both sequences only encoded by the K allele. Importantly, the K allele is the same allele associated with increased mammary *DGAT1* expression, which suggests a possible mechanism by which *DGAT1* K232A might exert its effect on *DGAT1* splicing, and hence mRNA expression.

The AAGAAG ESE motif has been proposed as the second-most common ESE hexamer in vertebrates [12]. Given the importance of ESEs for promoting splicing, we hypothesised that this motif could influence the splicing efficiency of *DGAT1* pre-mRNA to influence mRNA expression. It has been previously shown in humans that polymorphisms in ESEs can inhibit affinity for splicing factors and affect splicing, leading to altered mRNA and protein translation sequences that contribute to genetic disorders [13]. Additionally, the disruption of splicing has recently been reported for a novel *DGAT1* mutation in dairy cattle, whereby a non-synonymous A>C transversion in exon 16 disrupts a putative ESE motif and causes the skipping of this exon [14]. This polymorphism results in an enzymatically inactive DGAT1, which in the homozygous state results in a severe phenotype characterised by scouring and slow growth [14].

To investigate the hypothesis that the K232A polymorphism might influence *DGAT1* pre-mRNA processing, we defined a splicing efficiency phenotype to quantitatively measure splicing of intron 8 and other junctions of the gene. Association analysis using these splicing definitions revealed strong splice enhancement for five *DGAT1* introns, providing evidence supporting the mechanism by which this variant might influence mammary *DGAT1* mRNA expression. Critically, the mammary *DGAT1* intron splicing efficiency effects appeared to bear the same genetic signature underpinning the eQTL and milk production QTLs reported for this locus, that is, the association rankings for SNPs were similar for all QTLs. The direction of effects is also consistent with this hypothesis, where animals bearing the K allele have increased milk fat percentage, *DGAT1* expression and efficiency of splicing. Conversely, the animals bearing the A allele showed decreased *DGAT1* expression and splicing efficiency, as well as decreased milk fat percentage.

Splicing efficiency is dependent on a number of factors, with the likelihood of an intron being retained in mature mRNA reflecting the strength of the splice site, intron length, GC content, splicing factor expression and changes in chromatin structure [15]. As such, polymorphisms in ESEs and other splicing elements can influence transcription levels by modifying the strength of the recruitment of the splicing machinery to the junctions in the pre-mRNA transcript [16]. The splicing efficiency effect and increased *alternative* splicing for *DGAT1* suggest that there are a number of weak splice sites in the gene, and the presence of the ESE in the K allele enhances the recruitment of the splicing machinery to increase their usage, resulting in increased splicing of these junctions.

Interestingly, the junctions that had a splicing efficiency phenotype associated with K232A were distributed throughout the gene and included introns 1–3, and 11, which are several kb from the polymorphism and ESE motif. Similar to a recent study [17], the approach taken in this study accounted for any size bias and read coverage differences across the gene and subsequently revealed no relationship between the size of the intron and the splicing efficiency at the junction. The *DGAT1* intron 1 and 2 junctions, which contain the two largest introns, both exhibit a strong splicing efficiency effect, with the intron 2 junction exhibiting the most significant effect in this analysis. An explanation for why particular *DGAT1* junctions appear to be influenced by K232A genotype while others remain unaffected is unknown at this stage. It is possible that during pre-mRNA processing, the *DGAT1* junctions are processed in an order such that some junctions become rate-limiting steps in the process. If such bottlenecks exist, then the presence of the ESE could influence the efficiency of the processing of the intron 8 junction and the junctions that are subsequently processed. This would result in certain junctions exhibiting a splicing efficiency difference based on the presence or absence of the ESE, while the junctions prior to the bottleneck would remain unaffected. Ultimately, further experiments are required to understand the relationship between the activation of the exon 8 ESE in *DGAT1* and its influence on the splicing efficiency at multiple junctions in the gene.

Despite the strong association between *DGAT1* K232A and the expression and splicing efficiency phenotypes, there was still some possibility that one or more of these associations were due to LD effects exerted by an unknown *cis* regulatory variant. To more directly probe the function of K232A, mini-gene constructs were generated for the K and A alleles in the absence of native promoter sequence. Differing only by the dinucleotide substitution responsible for K232A, expression testing of these constructs replicated the splicing efficiency effect for a subset of the same junctions implicated *in vivo*, unequivocally assigning an expression-based mechanism to this variant.

Interestingly, the splicing efficiency effects appeared to result in an increase in spliced mRNA expression, rather than increased expression *per se* as there was no concomitant increase in unspliced transcripts at the two junctions exhibiting the splicing efficiency phenotype. The lack of increased expression of the unspliced pre-mRNA transcripts may be the result of an increased rate of pre-mRNA processing, and supports the hypothesis that splicing directly impacts mammary expression of mature *DGAT1* mRNA. A previous study [18] reported that many transcripts retain introns, and that these transcripts are retained in the nucleus without undergoing degradation via nonsense-mediated RNA decay. This study also showed that *in vitro*, these introns appear to be eventually spliced out at a much slower rate than other introns in the same transcript. These observations suggest that incompletely spliced *DGAT1* transcripts may also be spliced eventually, albeit at a slower rate.

While it is not the first time an expression-based effect of *DGAT1* has been proposed as the mechanism by which this gene influences milk composition [14], our study is the first to provide evidence supporting an expression-based effect associated with K232A. Association mapping at the *DGAT1* locus using imputed WGS variants showed that *DGAT1* K232A retains its status as one of the top ranking variants. However, K232A was not the marker with the smallest P-value, so the possibility remains that additional effects reside at the locus, or that imperfect sequence imputation or sampling error may have influenced the relative association rankings of the variants in this interval.

To attempt to address these possibilities, further association analysis was conducted to include K232A genotype as a covariate in the models. This analysis removed the majority of the association signal for *DGAT1* expression, suggesting that the *cis*-eQTL could be derived, for the most part, from *DGAT1* K232A. The clusters of highly significant markers in the previous analysis were no longer associated with *DGAT1* expression in these models, suggesting that these variants were tagging the signal from K232A. However, a seemingly distinct, statistically significant eQTL remained, signifying there may be additional effects on mammary *DGAT1* expression. A number of these highly associated markers are located upstream of the transcription start site of the gene, suggesting there may be an additional promoter driven effect on mammary *DGAT1* expression. One possibility is the previously-proposed VNTR polymorphism [5], which was hypothesised to increase the number of putative SP1 transcription factor binding sites, and stimulate an increase in *DGAT1* expression.

While we were able to convincingly demonstrate *cis*-eQTL and splicing efficiency effects at the *DGAT1* locus, an unresolved question is what proportion of the K232A impacts on milk composition are derived from differences in enzymatic activity and alternatively from the expression based effects. One possible option to delineate these two mechanisms would be to use redundant codons to create cell lines that encode identical DGAT1 proteins, yet have alternative ESE-encoding genomic sequences. Unfortunately, however, lysine and alanine amino acids have limited redundancy, precluding the design of such constructs.

## Summary and Conclusion

A QTL underpinned by the *DGAT1* gene represents one of the most well-known and validated bovine milk composition and production effects, presenting profound impacts on these traits. Despite the effects being long-attributed to an enzymatic mechanism as a consequence of a K232A missense mutation, we have used a large mammary RNAseq dataset in conjunction with *in vitro* expression experiments to highlight an alternative functional effect of this variant. Our experiments show that *DGAT1* is differentially expressed by QTL genotypes in the mammary gland, and confirm a splice enhancement role for the K232A mutation that potentially modulates these effects. Although the relative contribution of splice enhancement and differential enzymatic activities are unknown, these data suggest that the myriad lactation effects attributed to *DGAT1* K232A may, at least in part, derive from an expression-based mechanism.

## Acknowledgements

We gratefully acknowledge S. Morgan and staff at DairyNZ Ltd (Hamilton, New Zealand), and Phil McKinnon, Ali Cullum and staff at AgResearch (Hamilton, New Zealand) for enabling tissue sampling of lactating animals. We also acknowledge the services provided by AGRF, NZGL and the University of Auckland Centre for Genomics, Proteomics, and Metabolomics (Auckland, New Zealand) for RNA extraction and sequencing. Finally, we gratefully acknowledge the financial support provided by the Ministry for Primary Industries (MPI; Wellington, New Zealand), and Ministry of Business, Innovation and Employment (MBIE; Wellington, New Zealand), who independently co-funded aspects of the work through the Primary Growth Partnership, and Endeavour Fund research programs.

## Author Contributions

Conceived and designed experiments: TF, TL, MDL, RGS, RJS, SRD. Performed the experiments: TF, MDL, KT, TL, RH. Contributed analysis tools: KT, TL, TJ, RH. Analysed the data: TF, TL, MDL. Wrote the manuscript: TF, MDL, TL.

## Methods

### Ethics statement

All animal experiments were conducted in strict accordance with the rules and guidelines outlined in the New Zealand Animal Welfare Act 1999. For the mammary tissue biopsy experiment, samples were obtained in accordance with protocols approved by the Ruakura Animal Ethics Committee, Hamilton, New Zealand (approval AEC 12845). All other data were generated as part of routine commercial activities and were therefore outside the scope for formal committee assessment and ethical approval (as defined by the above guidelines). No animals were sacrificed for this study.

### DNA extraction and high throughput genotyping

The animals used for the analysis comprised 375 mostly Holstein-Friesian NZ dairy cows, representing a subset of 406 sequenced animals described in detail previously [10,19]. Briefly, genomic DNA was extracted from ear-punch tissue or blood by GeneSeek (Lincoln, NE, USA) and processed using Qiagen BioSprints kits (Qiagen).

Most (n=354) animals were genotyped with the Illumina BovineHD BeadChip by GeneSeek, with the remainder (n=21) genotyped on the Illumina BovineSNP50 panel, then imputed to the BovineHD panel as described previously [10,11]. A subset of 115 SNPs in the 1 Mbp interval centred on DGAT1 K232A (chr14:1302265–2302265) were assessed for this study. In addition to these markers, RNAseq-derived genotypes (see RNA sequencing section below) for K232A (chr14:1802265G>A SNP) were also included in the analyses, as this variant is the first base of the *DGAT1* K232A MNP. These genotypes were extracted using samtools [20] (version 0.1.19), with missing genotypes called by manual interrogation of RNAseq reads overlapping the variant. All genotypes were recoded using PLINK [21] (version 1.90b2c) to 0, 1 or 2 to represent the number of alternative alleles for each marker (i.e. 0, 1, and 2 to represent the homozygous reference, heterozygous, and homozygous alternative genotypes, respectively).

In addition to these genotypes, 3,128 imputed whole-genome sequence (WGS) derived variants in the 1 Mbp interval of interest were used. These were imputed with Beagle v4 [22], using a reference population of 556 animals as described previously [10,11].

### RNA sequencing

RNA sequencing and informatics was conducted prior to the experiments presented herein, and has also been described elsewhere [10,23]. Animals representing this dataset comprised three cohorts sampled at different points in time. Twenty-one of the samples were collected in 2004 and 2012 with sequencing conducted by NZGL (Dunedin, New Zealand) using the Illumina HiSeq 2000 instrument. For these samples, libraries were prepared using the TruSeq RNA Sample Prep Kit v2 (Illumina). RNA sequencing of two batches of samples (183 and 171 samples respectively) collected in 2013 was carried out by the Australian Genome Research Facility (AGRF; Melbourne, Australia) using the Illumina HiSeq 2000 instrument. Libraries for these two cohorts were prepared using the TruSeq Stranded Total RNA Sample Prep Kit (Illumina) with ribosomal depletion using Human/Mouse/Rat Ribo-Zero kit (Epicentre/Illumina). All samples were sequenced using a 100 base-pair (bp) paired-end protocol, with two samples multiplexed per lane.

RNA sequence data representing the 375 animals were mapped to the UMD3.1 genome using Tophat2 [24] (version 2.0.12), locating an average of 88.9 million read-pairs per sample. Cufflinks software [25] (version 2.1.1) was used to quantify expressed transcripts, and yielded fragments per kilobase of exon model per million mapped (FPKM) expression values. In addition, read counts were determined using HTseq software [26] (version 0.6) and processed using the variance-stabilising transformation (VST) normalisation method in DESeq [27] (version 1.18) to derive gene expression phenotypes suitable for linear model analysis, and subsequent eQTL mapping.

To investigate the influence of *DGAT1* K232A on *DGAT1* splicing efficiency, the number of reads mapping to each intron and exon of *DGAT1* was determined using HTSeq 0.6.0 [26], with the intron and exon boundaries specified by the RefSeq annotation (NM_174693.2). The splicing efficiency phenotype for *DGAT1* intron 8 was calculated as the percentage of *DGAT1* RNAseq reads mapping to the intron. The splicing efficiency phenotypes for each individual RefSeq *DGAT1* junction were calculated as the ratio of exonic reads to intronic reads corresponding to the junction (of spliced and unspliced reads, respectively). Reads were considered exonic if they bridged the splicing junction i.e. mapped to the 3′ end of the preceding exon and the 5′ end of the following exon. Reads were considered intronic or unspliced if they mapped to the 3′ end of the preceding exon and through the intron-exon boundary into the intron.

### Genetic association analysis

Associations between K232A and the 115 Bovine HD SNPs in the 1 Mbp interval surrounding *DGAT1* K232A and *DGAT1* expression were quantified using pedigree-based mixed models in ASReml-R [28,29]. Additionally, associations between the 3,128 imputed WGS-derived variants in the 1 Mbp interval centred on *DGAT1* K232A and *DGAT1* expression were quantified in the same way. Each SNP was fitted in a separate sire-maternal grandsire single trait model, with SNP treated as a quantitative variable based on the number of copies of the alternative allele, and variance components estimated in a restricted maximum-likelihood (REML) framework. Covariates for sequencing cohort, the proportions of NZ Holstein-Friesian ancestry, US Holstein-Friesian ancestry, Jersey ancestry and heterosis effects were also included in the models.

The additive genetic variance for each SNP was calculated using 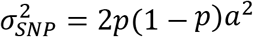, where *p* is the frequency of the highest frequency allele and *a* is the estimated allele substitution effect. Polygenic genetic variances were evaluated as 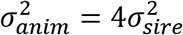 where 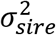 is the estimate of sire variance from the model. Total genetic variance was evaluated as 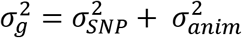 and phenotypic variance was evaluated as 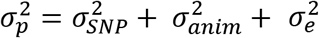 where 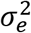 is the residual variance. The proportion of phenotypic variance explained by each SNP for each phenotype was calculated as 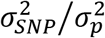 and the proportion of genetic variance explained by each SNP was calculated as 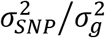.

Association analyses for splicing efficiency were also conducted using ASReml-R as described above. This analysis fitted 115 BovineHD SNPs plus K232A genotypes derived from the RNAseq data, with phenotypes defined as described in the RNA sequencing methods.

The Bioconductor package DEXSeq [8] was used to investigate the alternative splicing of *DGAT1* exon 8, which has previously been demonstrated to be associated with the *DGAT1* K232A genotype [4]. The genetic coordinates for the alternative form of *DGAT1* exon 8 (chr14:1802251–1802259 alternative, 1802260–1802325 reference) were manually added to the Enseml gene transfer format (GTF) file that contained gene structures of all the genes in the reference genome. Then, DEXSeq was used to count the relative usage of these two bins. A single factor ANOVA was used to test for the influence of the K232A genotype on the alternative splicing of DGAT1 exon 8.

### *In vitro* splicing efficiency assay

To test the effect of the K232A on *DGAT1* splicing efficiency *in vitro*, MAC-T cells [30] were transfected with *DGAT1* mini-gene constructs containing either the K232 or the A232 allele. The *DGAT1* alleles were based on the reference sequence (Accession number AY065621), and were identical with the exception of the AA>GC MNP that causes the K232A amino acid substitution. The *DGAT1* 5′ UTR was extended by 84 bp to represent the UTR apparent from mammary RNAseq data, and the first two introns were removed due to constraints on total insert size. Intron 1 is 3,616 bp and intron 2 is 1,943 bp, such that the collective 5,559 bp from these two introns is larger than the rest of the gene structure combined (3,117 bp; Supplementary Figure 1). Sequences representing the two *DGAT1* isoforms were synthesised and cloned into pcDNA3.1 by GenScript (New Jersey, USA). Co-transfection of cells with pMAXGFP plasmid (Lonza) was conducted in a 1:1 ratio to provide a normalisation control for transfection efficiency.

Cells were plated in 24-well plates and grown for 24 hours in proliferation media to achieve approximately 70% confluency. For cell transfection, 0.5 μL Lipofectamine® LTX (Invitrogen) was gently mixed with 25 μL Opti-MEM reduced serum media (Invitrogen). Aliquots containing 375 ng of both DGAT1 pcDNA3.1 constructs and pMAXGFP plasmid DNA, and 0.5 μL PLUS reagent were diluted in 25 μL Opti-MEM. The diluted plasmids were combined with the Lipofectamine® LTX, gently mixed and incubated at room temperature for 5 minutes, after which 50 μL transfection mix was added to each well. After 24 hours of incubation at 37°C, the cells were visualised on a Nikon TiE inverted light microscope prior to RNA extraction. All experiments were repeated in triplicate in three separate cell preparations from MAC-T passage numbers 9, 10, 11, and 12.

RNA was extracted from each well of a 24-well plate using a TRIzol-based protocol and was subjected to two sequential DNase treatments before quantification. Following DNase treatment, cDNA synthesis was performed using 2.5 μg of RNA as input for each 20 μL reaction. Complementary DNA (cDNA) was diluted 1:10 in Ultra-Pure water (Invitrogen) and used immediately for qPCR or stored at −20°C. Serial 5x cDNA dilutions were used to generate standard curves for each real-time PCR assay by pooling 4 μL from each experimental sample. Real-time PCR reactions were carried out in 10 μL volumes in 384-well plate format using standard qPCR cycling conditions for use with the LightCycler480^®^ Universal Probe System. Eukaryotic translation initiation factor 3K (*EIF3K*) was used as an endogenous control gene for normalisation of gene expression [31]. In addition, an assay was designed for the pMAXGFP plasmid as a further control to normalise for transfection efficiency.

To quantify splicing efficiency at the intron-exon junctions in *DGAT1*, assays were designed using Universal Probe Library and Primer 3 software to generate two assays for each junction. These assays were designed such that they had a common primer (either forward or reverse), and used the same probe. The expression of spliced mRNA transcripts was measured using primers that bound to the two exons adjacent to the intron/exon junction. The expression of unspliced mRNA transcripts was measured using a primer that bound to one of the adjacent exons and a primer that bound across the intron/exon junction. The average expression of each transcript across triplicate wells was calculated relative to the geometric mean of expression for the reference gene assays for each sample. Supplementary Figure 2 represents the amplicon design strategy used to differentiate spliced and unspliced *DGAT1* transcript forms. Supplementary Table 1 lists the primers and probes used for these experiments.

The average expression of the spliced transcripts was divided by the unspliced transcripts to get the splicing ratio for each junction for each sample. Student’s t-test was used to determine the statistical significance of differences between the two alleles, comparing the splicing ratio or mean spliced or unspliced transcript expression values for each junction.

---

**Supplementary Figure 1:**
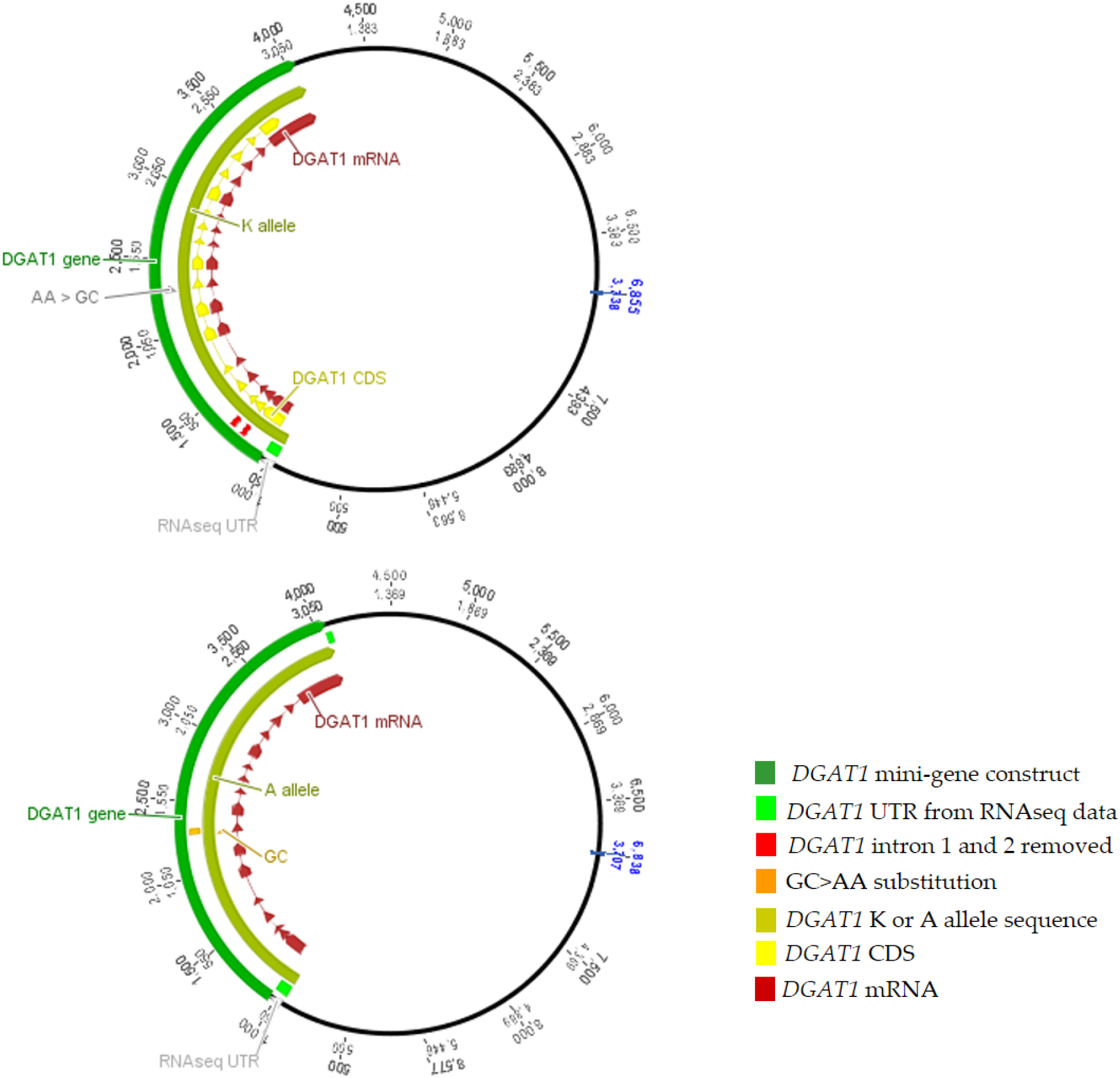
Schematic of the two *DGAT1* constructs inserted into pcDNA3.1.

**Supplementary Figure 2:**
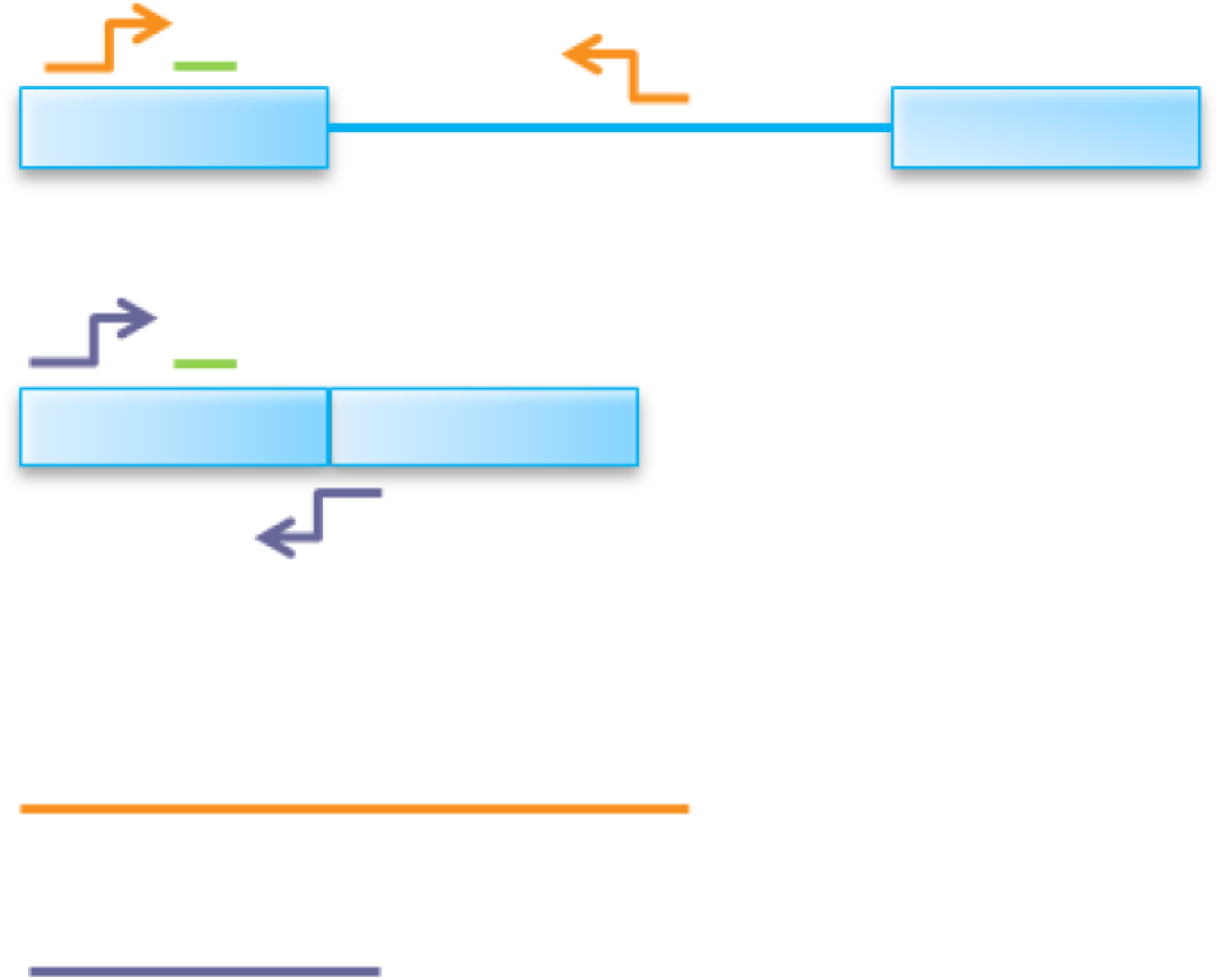
Schematic of the two RT-qPCR assays for each junction in *DGAT1*. The blue boxes represent exons while the blue line represents the intron. The green line represents the probe, while the orange and purple arrows represent the primers for unspliced and spliced mRNA transcripts, respectively. The first assay quantifies the intron containing pre-mRNA transcripts (orange) while the second assay quantifies the spliced mRNA transcripts (purple). The ratio of mRNA:pre-mRNA transcripts is used to generate a splicing efficiency phenotype for each junction.

**Supplementary Table 2:**
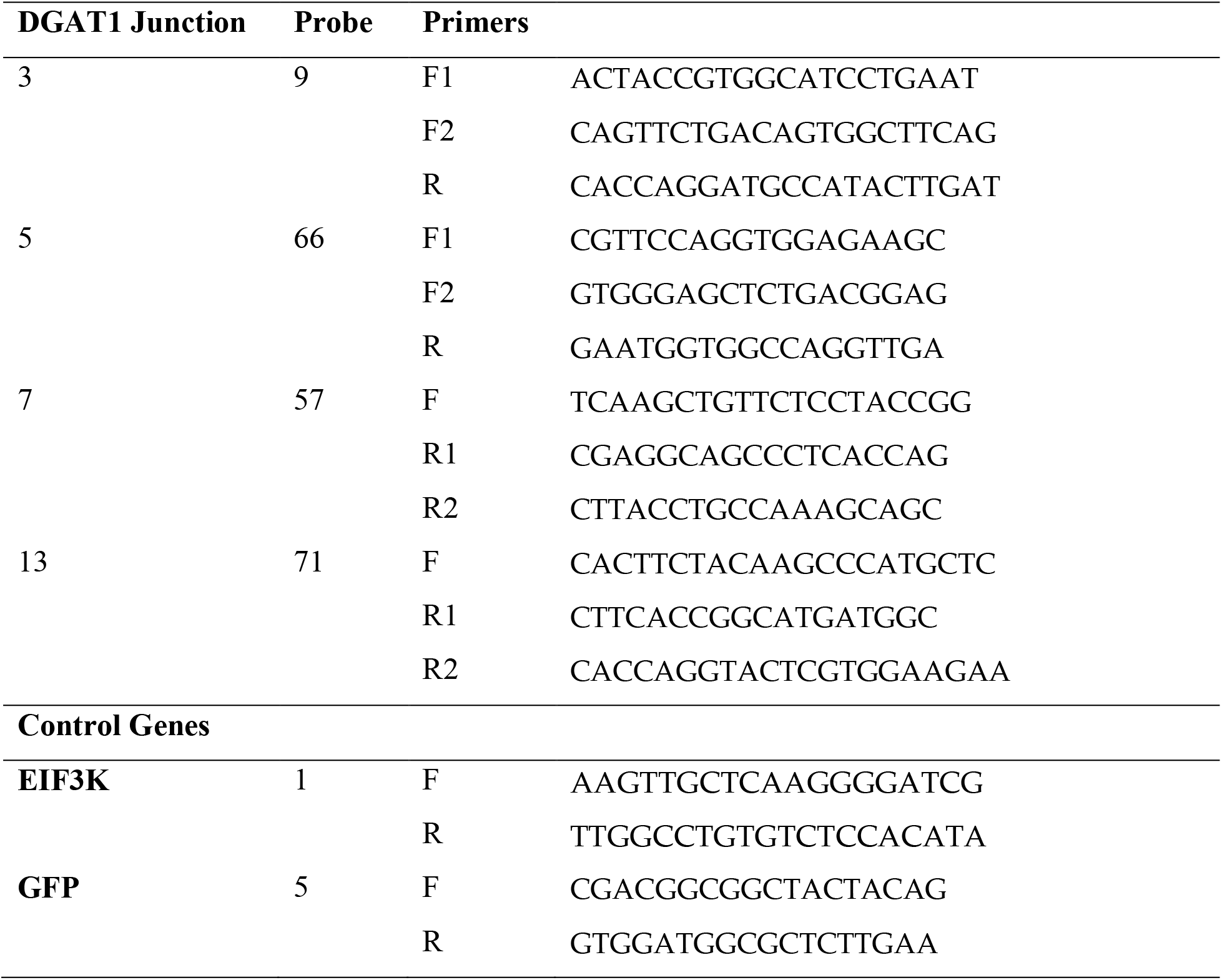
Primer sequences and assay design for RT-qPCR of *DGAT1* intron 3, 5, 7, and 13 junctions.

